# Mitochondrial active Ras2 protein promotes apoptosis and regulated cell death in a cAMP/PKA pathway-dependent manner in budding yeast

**DOI:** 10.1101/2021.10.13.464237

**Authors:** Barbara Bonomelli, Enzo Martegani, Sonia Colombo

## Abstract

In previous papers, using the eGFP-RBD3 probe, which binds Ras-GTP with high affinity, we showed that activated Ras proteins are localized to the plasma membrane and in the nucleus in wild-type *Saccharomyces cerevisiae* cells growing exponentially on glucose, while an aberrant accumulation of activated Ras in mitochondria correlates to mitochondrial dysfunction, accumulation of ROS and an increase of apoptosis. In this paper, we show that lack of *TPS1*, which is known to trigger apoptosis in *S. cerevisiae*, induces localization of active Ras proteins in mitochondria, confirming the above-mentioned correlation. Next, by characterizing the *ras1*Δ and *ras2*Δ mutants concerning localization of active Ras proteins and propensity to undergo cell death, we show that active Ras2 proteins, which accumulate in the mitochondria following addition of acetic acid, a well-known pro-apoptotic stimulus, might be the GTPases involved in regulated cell death, while active Ras1 proteins, constitutively localized in mitochondria, might be involved in a pro-survival molecular machinery. Finally, by characterizing the *gpa2*Δ and *cyr1*Δ mutants concerning the propensity to undergo cell death, we show that active mitochondrial Ras proteins promote apoptosis through the cAMP/PKA pathway.

## 1. Introduction

The regulated cell death (RCD) includes a set of mechanisms aimed at the targeted elimination of superfluous, irreversibly damaged and/or potentially harmful cells, with an advantage for the body’s homeostasis. RCD, as well as autophagy, can be activated in response to cellular stress. Dysfunctions in the regulation of RCD underlie several human diseases [1]. RCD is evolutionarily conserved among mammals and unicellular eukaryotes, such as the yeast *Saccharomyces cerevisiae*, which is one of the model organisms used to study apoptosis, a form of regulated cell death [2, 3]. Several yeast genes involved in apoptosis have been identified, presenting an ortholog in mammals. These include genes encoding a caspase (Yca1), apoptosis inducing factor (Aif1), OMI serine protease (Nma111) and endonuclease G (Nuc1) [4–8]. The apoptotic pathway can be activated by the expression of pro-apoptotic heterologous proteins such as Bax [9], but also by exogenous agents such as hydrogen peroxide and acetic acid [10, 11]. In particular acetic acid has been widely used as inducer of apoptosis in yeast [5, 11], since according to Sokolov et al [12] it triggers a ROS (Reactive Oxygen Species) dependent death.

The *Saccharomyces cerevisiae* contains two RAS genes, *RAS1* and *RAS2* coding for two proteins showing a significant homology to the mammalian Ras proteins [13, 14]. Yeast Ras1 and Ras2 proteins, like their mammalian counterparts, undergo extensive posttranslational modification (farnesylation, palmitoylation and carboxymethylation) and are deposited on the inner surface of the plasma membrane by a specialized transport mechanism [15–17]. The Ras proteins belong to a superfamily of small GTPases that cycle between an inactive state, when loaded with GDP, and an active state, when loaded to GTP. In budding yeast, the Ras proteins activity is tuned by two classes of regulatory proteins: Guanine nucleotide Exchange Factors (GEFs) Cdc25 and Sdc25, which stimulate the GDP/GTP exchange and Ira1 and Ira2 GTPase Activating Proteins (GAPs), which stimulate the intrinsically low Ras GTPase activity [18]. The molecular mechanisms of modulation of Ras proteins activity have been conserved during evolution, as it was confirmed by the functional interchangeability of Ras proteins and their regulators in mammals and budding yeast [18, 19].

In previous papers [20–22] we give evidences that in budding yeast exists a correlation between mitochondrial localization of Ras-GTP and apoptosis. In particular, we demonstrated that deletion of *WHI2*, a gene coding for a protein known to influence cell cycle exit under conditions of nutritional stress, leads to the loss of coordination between nutritional sensing and actin regulation, resulting in the failure to correctly traffic Ras2 to the vacuole. Consequently, Ras2 protein localizes to the mitochondrial surface in its active form. This leads to a failure to shut down Ras signalling, to mitochondrial dysfunction, accumulation of damaging ROS and cell death [20] In a subsequent work, we showed that addition of acetic acid to *S. cerevisiae* wild type cells causes within five minutes a delocalization of active Ras from plasma membrane and nucleus to mitochondria [21]. Furthermore, we demonstrated that addition of either acetic acid or hydrogen peroxide to *hxk2* cells, showing a constitutive localization of active Ras at the mitochondria, causes an increase in the level of ROS, mitochondrial dysfunctions and an increase of both apoptotic and necrotic cells compared with the wild-type strain [21, 22]. Finally, we showed that also lack of *SNF1*, the homolog of the AMP-activated protein kinase (AMPK) in *S. cerevisiae*, induces localization of active Ras in mitochondria and triggers apoptosis in this microorganism [23].

The two Ras proteins of *S*.*cerevisiae*, Ras1 and Ras2 cannot be discriminated by the eGFP-RBD3 probe we used to investigate the localization of active Ras proteins (Ras-GTP) [24]. Consequently, in order to explore which one of these two GTPases is actually involved in apoptosis and cell death, we characterized the *ras1*Δ and *ras2*Δ mutants concerning localization of active Ras proteins and propensity to undergo these cellular *processes*, following a pro-apoptotic stimulus. We show that the Ras2 protein might be the GTPase involved in apoptosis and cell death induced by addition of acetic acid, while active Ras1 protein is localized in mitochondria and might be involved in a pro-survival molecular machinery. Next, we aimed to investigate whether the accumulation of active Ras proteins in the mitochondria determines an increase in the level of ROS, apoptosis and cell death in a cAMP/PKA pathway dependent manner [25, 26], since data reported in literature show that many proteins of the Ras/cAMP pathway, including Ira2, Cyr1, Bcy1 and Tpk1 are located in the mitochondria [27,28], suggesting that Ras-dependent transduction pathways could also originate and/or be “managed” by these organelles. Our results show that indeed active mitochondrial Ras2 protein promotes apoptosis and cell death through the cAMP/PKA pathway. Finally, we show that also in the *tps1*Δ mutant [29, 30] a correlation exists between aberrant accumulation of activated Ras in mitochondria and propensity of the cells to undergo apoptosis.

## 2. Materials and Methods

### 2.1. Yeast strains, plasmids and media

The yeast strains used in this study are indicated in Table 1.

**Table 1.** List of yeast strain

| Strain | Main Genotype | Source/reference |
| --- | --- | --- |
| W303-1A | *MATa ade2-1 can1-100 his3-11,15 leu2-3112 trp1-1 ura3-1* | Thomas and Rothstein, 1989 [31] |
| YSH290 | W303-1A with *tps1::TRP1* | Hohmann et al., 1993 [32] |
| SP1 | *MATa his3 leu2 ura3 trp1 ade8 can1* | Nikawa *et al.* [33] |
| ST-1 | SP1 with *ras1::URA3* | kindly provided by J. Thevelein KU Leuven [34] |
| KP-2 | SP1 with *ras2::URA3* | kindly provided by J.Thevelein, KU Leuven [34] |
| SC591 | W303-1A with *ras2::URA3* | Our laboratory |
| W303-1A *gpa2Δ* | W303-1A with *gpa2::LEU2* | kindly provided by J. Winderickx., KU Leuven |
| GG104 | W303-1A with *pde2::TRP1*, *cyr1::KanMX2*, *msn2::HIS3*, *msn4::TRP1* | Roosen et al., (2005) [35] |

The plasmid pYX212-eGFP-RBD3 encoding for the eGFP-RBD3 probe [24] was used to localize Ras-GTP in yeast cells through fluorescence microscopy analyses. Deletion of *RAS2* in the W303-1A strain was made using polymerase chain reaction (PCR)-based strategy as previously described [36]. In particular, a PCR-cassette containing *URA3* amplified from plasmid YCp50 was used. All the primers used in this study are listed in Supplementary Table I. Yeast cells were transformed according to Gietz et al. [37]. Rich medium contained 2% w v^-1^ glucose, 2% w v^-1^ peptone, 1% w v^-1^ yeast extract (YPD). Synthetic complete media (SD) contained either 2% glucose or 2% galactose, 6.7 g/l YNB w/o amino acids (supplied by ForMedium^™^, United Kingdom) and the proper selective drop-out CSM (Complete Synthetic Medium, supplied by ForMedium^™^, United Kingdom). Culture density was measured with a Coulter Counter (Coulter mod. Z2) on mildly sonicated, diluted samples and we monitored the growth and defined the exponential phase by measuring the cell number/ml at discrete interval (30 min) and plotting the log of cell concentration as a function of time. YEPD plates contained 2% w v^-1^ glucose, 2% w v^-1^ peptone, 1% w v^-1^ yeast extract and 2% w v^-1^ agar.

### 2.2. Acetic acid treatment

Cells were grown at 30° C to exponential-phase (in the range 0.6-1 × 10^7^ cells mL^-1^) in SD medium, harvested, resuspended (on average 10^7^ cells mL^-1^) in fresh SD medium adjusted to pH 3.0 (set with HCl) and treated with acetic acid (Riedel-deHaeden) at the indicated concentration. Cells were incubated for 200 min at 30° C with shaking (160 rpm) [21].

### 2.3. Viability assay

Cells were grown in SD medium at 30° C until exponential phase (in the range 0.6-1× 10^7^ cells mL^-1^) and treated with 80 mM acetic acid for 200 min. Cell number was measured before and after acetic acid treatment using a Coulter Counter (Coulter mod. Z2) and 1000 cells were plated on YEPD agar plates in triplicate. Viability was determined by measuring colony forming units (cfu) after 3 days of growth at 30° C [21].

### 2.4. Dihydrorhodamine 123 (DHR123) staining

ROS were detected with DHR123 (Sigma Aldrich) essentially as described by Madeo et al. [10, 21]. Cells were grown in SD medium at 30° C until exponential phase (in the range 0.6 × 10^7^ cells mL^-1^) and treated with acetic acid as described above for 80 min. Then, DHR123 was added directly to the culture medium at the final concentration of 5 µg mL^-1^ (from a 2.5 µg µL^-1^ stock solution) and incubation at 30° C with shaking was prolonged for additional 120 min in the dark at 30° C. After a total of 200 min incubation, cells were diluted in 50 mM filtered TrisHCl pH 7.5 to 10^6^ mL^-1^ and analysed using a cytofluorimeter (CytoflexS, Beckman) with excitation and emission wavelength of 488 and 525 nm respectively. A total of 20.000 events were acquired for each sample and data were processed using CytExpert software.

### 2.5. Annexin V and propidium iodide (PI) staining

(FITC)-conjugated recombinant Annexin V (Immuno Tools) was used for the detection of phosphatidylserine exposed in the membrane of apoptotic cells as reported previously [21–23]. Cells were harvested after 200 min of acetic acid treatment, washed with sorbitol buffer (1M sorbitol, 0.1M NaH2PO4, pH 8.0) and the cell wall was digested with Zymolyase 20T (Seikagaku Biobusiness Corporation) for about 35 min at 37° C. Cells were then washed two times with binding buffer (10 mM Hepes/NaOH pH 7.4, 140 mM NaCl, 2.5 mM CaCl2, 1.2 M sorbitol). Spheroplasts were resuspended in 35 µL of binding buffer and incubated with 2.5 µL of Annexin V (ImmunoTools) and 2 µL of a PI (Fluka) working solution (50 µg mL^-1^ in 10 mM TrisHCl pH 7.0) for 15 min in the dark at room temperature. After staining, the samples were resuspended in binding buffer and analysed using a cytofluorimeter (CytoflexS, Beckman) with excitation at 488 nm and 525 nm emission for Annexin and 585 nm emission for PI. A total of 30.000 events were acquired for each sample and data were processed using CyExpert software.

### 2.6. Fluorescence microscopy

Cells expressing the eGFP-RBD3 probe [24] were grown in SD medium at 30° C until exponential phase(in the range 0.6-1 × 10^7^ cells mL^-1^) and incubated with the mitochondrial marker rhodamine B hexyl ester perchlorate (Molecular Probes, Eugene, OR) 100 nM final concentration for about 5 min before imaging. Images were acquired with a Nikon Eclipse 90i microscope equipped with a 60X oil immersion objective and a standard FITC filter set for GFP fluorescence. Images were recorded digitally using a Nikon DS-Qi1Mc camera and processed using Adobe Photoshop (Adobe Systems, Inc.) and ImageJ software (https://imagej.nih.gov). Determination of the spatio-temporal localization of Ras-GTP after addition of glucose to *tps1Δ* cells growing exponentially on 2% galactose medium was performed as following. Cells were grown in medium containing 2% galactose at 30° C till exponential phase. Subsequently, 100 µl of cells were seeded on concanavalin A-coated cover glass for 10 min. The cover glass was washed four times using 1 ml of 2% galactose medium and put on top of a Thoma chamber. Images were acquired before and after addition of glucose with a Nikon Eclipse 90i microscope equipped with a 60X oil immersion objective and a standard FITC filter set for GFP fluorescence. Glucose was added by pipetting, between the cover glass and the Thoma Chamber, 45 µl of 4% glucose medium. To avoid bleaching, the fluorescence images were acquired for 4s every 1 minute for 10 min and the shutter was kept off in the meantime. The time series of images was then processed using ImageJ software.

### 2.7. Statistical analysis

All the experiments were done at least in triplicate and the mean and Standard deviation was shown. The Student’s t test was used for assessing the significance of the experimental data. The experimental data were elaborated with Excel TM.

## 3. Results and discussion

### 3.1. Lack of TPS1 gene induces localization of active Ras in mitochondria in budding yeast

In the yeast S*accharomyces cerevisiae*, trehalose-6P-synthase (Tps1) is involved in trehalose synthesis, a highly stable disaccharide, implicated in many cellular processes and especially known for its protective role in response to a variety of environmental and nutritional stresses [38]. In particular, Tps1 first produces trehalose-6-phosphate (T6P) from glucose-6P and UDP-Glucose, which is then dephosphorylated by a specific trehalose-6P-phosphatase (Tps2p), to yield to trehalose. Recent data published in literature show a new function for the Tps1 protein as a pro-survival factor during growth and apoptotic stress [39,40]. In particular Peeters et al [40] showed that addition of glucose to *tps1*Δ cells grown on galactose triggers activation of Ras proteins and apoptosis. In previous papers, we show that activated Ras proteins (Ras-GTP) are localized to the plasma membrane and in the nucleus in wild-type yeast cells growing exponentially on glucose [24], while an aberrant accumulation of activated Ras in mitochondria correlates to mitochondrial dysfunction, accumulation of ROS and an increased sensitivity to pro-apoptotic treatments [20–23]. Therefore, in this paper we investigated the localization of active Ras proteins in the *tps1*Δ strain using the eGFP-RBD3 probe [24] in order to verify whether also in this mutant a correlation exists between aberrant accumulation of activated Ras in mitochondria and apoptosis. Due to the inability of the *tps1*Δ mutant to grow on glucose [29, 32], we performed the experiment during growth on galactose medium. As a control, the localization of active Ras proteins was also investigated in W303-1A cells during growth on the same medium. As expected and in agreement with previously published data [24], in W303-1A cells growing on medium containing galactose, active Ras proteins were partially localized to the plasma membrane, in the nucleus and in mitochondria. As shown in Figure 1, in *tps1*Δ cells active Ras proteins were almost totally localized in mitochondria (90%), indicating that also in this mutant a correlation exists between aberrant accumulation of activated Ras in mitochondria and propensity of the cells to undergo apoptosis. Moreover, we show that glucose addition to galactose-grown *tps1*Δ cells triggered a fast and marked increase of green fluorescence in mitochondria, suggesting that there is an increase of active Ras in these organelles (Figure 2). This result agrees with published data showing that glucose addition to galactose-grown *tps1*Δ cells triggers a rapid and dramatic activation of Ras [40].

**Figure 1.**
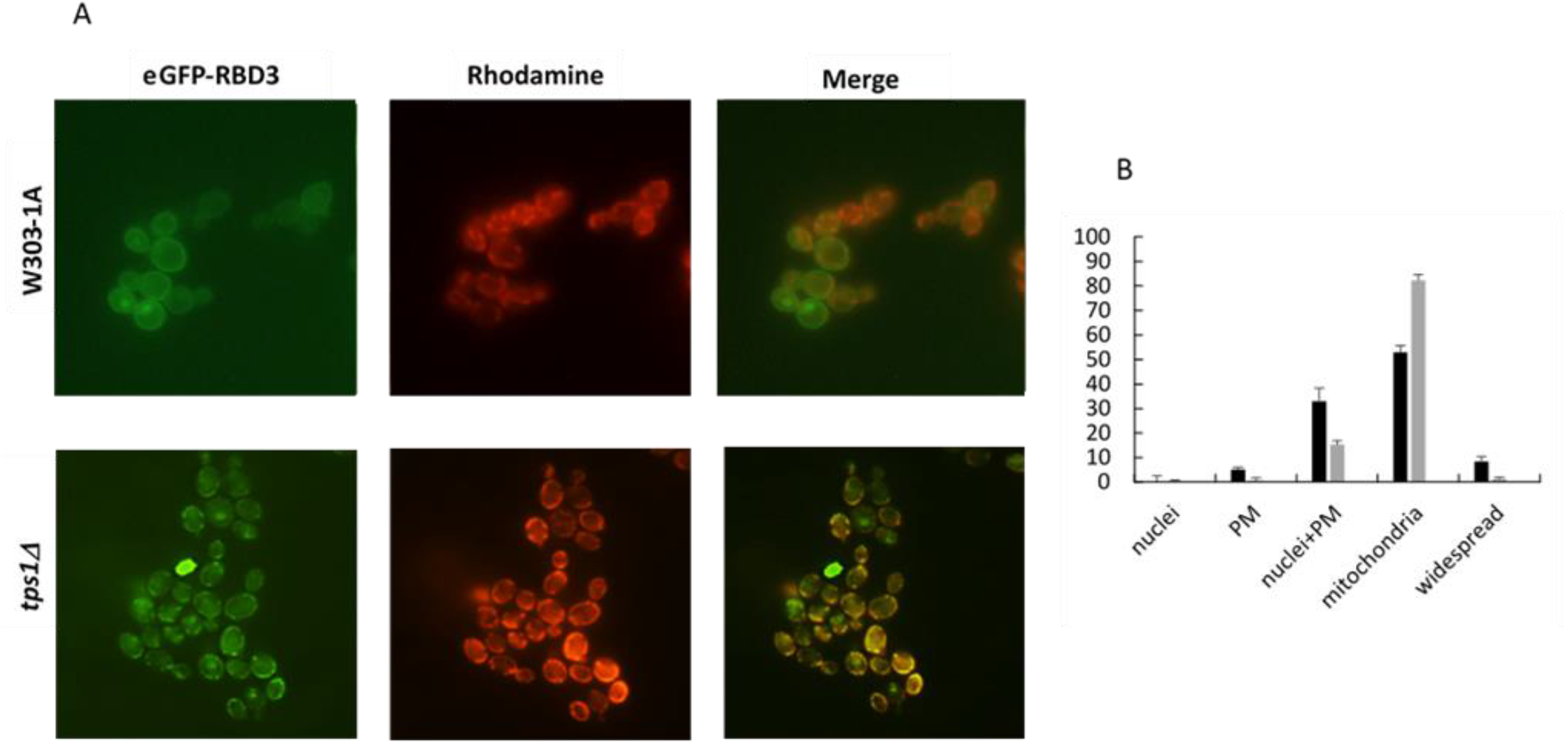
Localization of active Ras proteins in the tps1 Δ mutant. (A) Localization of eGFP fluorescence and red-fluorescent rhodamine B hexyl ester in W303-1A and *tps1*Δ cells expressing the eGFP-RBD3 probe and growing exponentially on galactose medium. (B) Subcellular distribution of eGFP fluorescence in W303-1A (black) and *tps1*Δ (grey) cells. Cells belonging to each class were counted and expressed as a percentage on the total number of fluorescent cells. Approximately 300 cells were scored and error bars represent the standard deviation.

**Figure 2.**
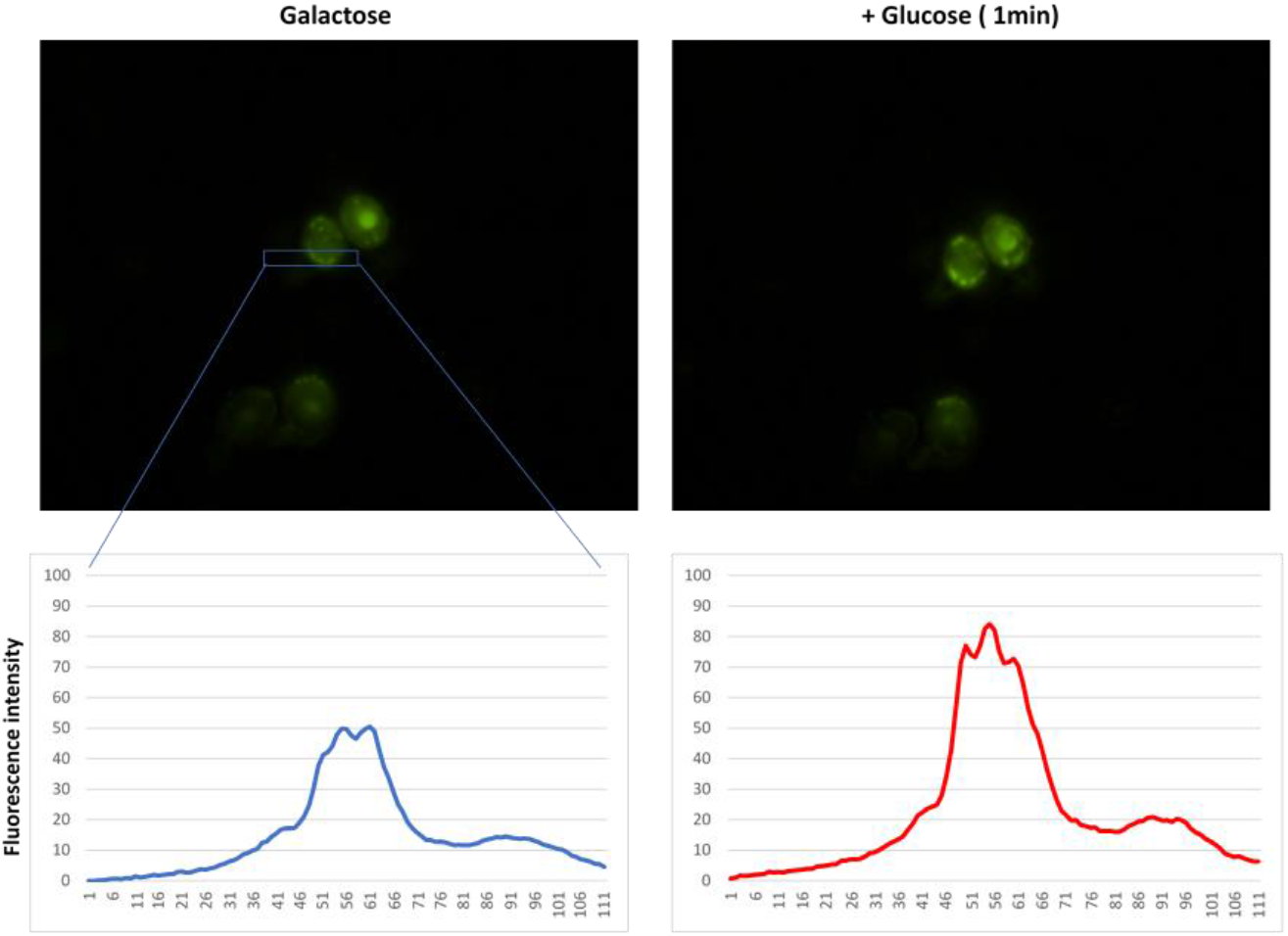
Glucose triggers a rapid and marked increase of fluorescence in mitochondria of tps1Δ cells. eGFP fluorescence level before and 1 min after addition of glucose to *tps1*Δ cells expressing the eGFP-RBD3 probe and growing on galactose medium. Cells growing in 2% galactose medium were seeded on concanavalin A-coated cover glass and mounted on top of a Thoma chamber (see Materials and Methods for details). Images were acquired before and 1 minute after addition of glucose with a Nikon Eclipse 90i microscope equipped with a 60X oil immersion objective and with the same setting of exposure time and imaging (4s, 1.2 gain) (upper). Images was than analysed with ImageJ software to obtain the pixel values (profile) (bottom) along the rectangular ROI overimposed in blue in the figure.

### 3.2. The Ras2 protein is the GTPase involved in apoptosis induced by addition of acetic acid

*S. cerevisiae* has two Ras proteins, Ras1 and Ras2 [13]. Since the eGFP-RBD3 probe is unable to discriminate between them, in order to investigate which one of the two proteins was involved in apoptosis, we characterized the *ras1*Δ and *ras2*Δ mutants concerning localization of active Ras proteins and propensity to undergo apoptosis and cell death. Our results show that in glucose growing cells, deletion of *RAS2* caused a mitochondrial localization of the probe, while deletion of *RAS1* did not impair the nuclear and plasma membrane localization of active Ras2 (Figure 3). We also investigated the localization of active Ras2 protein after addition of 40 mM acetic acid, a well-known inducer of apoptosis in yeast, to *ras1*Δ cells. We could show that in this strain, active Ras2 protein moved from plasma membrane and nuclei to the mitochondria within 5 minutes, suggesting that active Ras2 proteins might actually be the GTPases involved in apoptosis and cell death (Figure 3B). However, a possible contribution of active Ras1 protein in this cellular process might not be excluded. In order to investigate the role of active Ras1 proteins in apoptosis and cell death, we conducted cell survival experiments, assessed oxidative damage (ROS level) and the expression of cytological markers of apoptosis and necrosis, in the presence and absence of acid acetic, in the *ras2*Δ strain.

**Figure 3.**
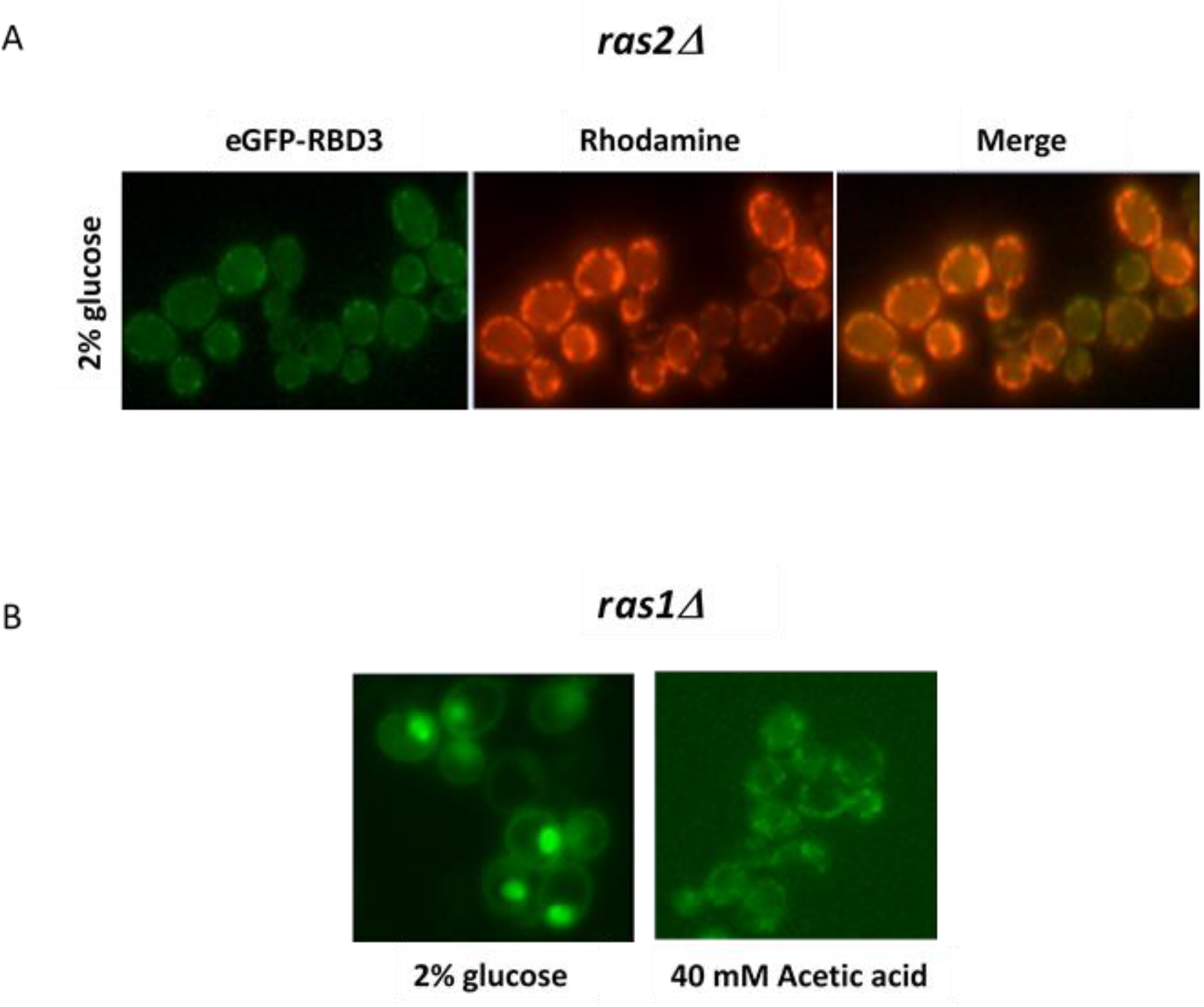
Localization of active Ras proteins in RAS deletion mutants. (A) Localization of eGFP fluorescence and red-fluorescent rhodamine B hexyl ester in SP1 *ras2*Δ cells expressing the eGFP-RBD3 probe and growing exponentially on glucose medium. (B) Localization of eGFP fluorescence in SP1 *ras1*Δ cells expressng the eGFP-RBD3 probe, growing exponentially on glucose medium and 5 minutes after addition of 40 mM acetic acid.

After treatment with 80 mM acetic acid for 200 minutes at 30°C, *ras2*Δ cells showed a significant increase in cell survival when compared with wild-type cells (Figure 4A). Congruently, the amount of ROS was significantly reduced in *ras2*Δ cells compared to wild-type cells (Figure 4B, Supplementary Figure 2). We also measured the level of cytological markers of apoptosis and necrosis in the r*as2*Δ mutant by flow-cytometry. In particular, combined Annexin V/propidium iodide (PI) staining was used to discriminate between early apoptotic (Annexin V+/PI-), late apoptotic/secondary necrotic (Annexin V+/PI+) and primary necrotic (Annexin V-/PI+) deaths. Our results showed that the percentage of *ras2*Δ cells in late apoptosis/secondary necrosis following treatment with 80 mM acetic acid was significantly reduced compared to that obtained for the corresponding wild strain (Figure 4C (c), Supplementary Figure 3), while the percentage of primary necrotic cells was very low, both in *ras2*Δ and wild type strains (Figure 4C (b)). In summary, data on cell survival, ROS level and percentage of cells in late apoptosis/secondary necrosis are consistent and indicate that constitutively mitochondrial localized active Ras1 protein is not related to the propensity of the cells to undergo a regulated cell death modality with characteristic features of apoptosis, following addition of acetic acid. However, unexpectedly, the percentage of cells in early apoptosis in the *ras2*Δ strain, following treatment with acetic acid, was significantly high compared with that obtained for the wild strain (Figure 4C (a)). This data, which is apparently in contrast with the above data, might indicate that in yeast, as well as in higher eukaryotes, apoptosis is divided into phases, being the initial phase (early apoptosis) reversible and secondary necrosis (combined necrotic and apoptotic features) the final stage of the apoptotic process [3]. Moreover, the presented results are congruent with recent data demonstrating that late apoptosis/secondary necrosis might not be necessarily an accidental consequence of early apoptosis, but a finely regulated process controlled by a specific molecular machinery [41] and that secondary necrotic cells might actually undergo secondary necrosis following other cell death subroutines [3]. In conclusion, taken together our data suggest that: 1) active Ras2 proteins, which accumulate in the mitochondria following a pro-apoptotic stimulus (acetic acid for instance), might be the GTPases involved in apoptosis and cell death; 2) mitochondrial active Ras1 proteins might be involved in a pro-survival molecular machinery. The two proteins Ras1 and Ras2 have similar and overlapping function in the activation of adenylate cyclase [42, 43], but they have different expression at the level of mRNA and protein [44]. In addition, Ras2 is much more abundant than Ras1 [44, 45] (20000 mol/cell versus 2000 of Ras1 [45]); moreover, it has been shown that these two proteins have an opposite function also in replicative aging of yeast cells [46] and in stress response [47].

**Figure 4.**
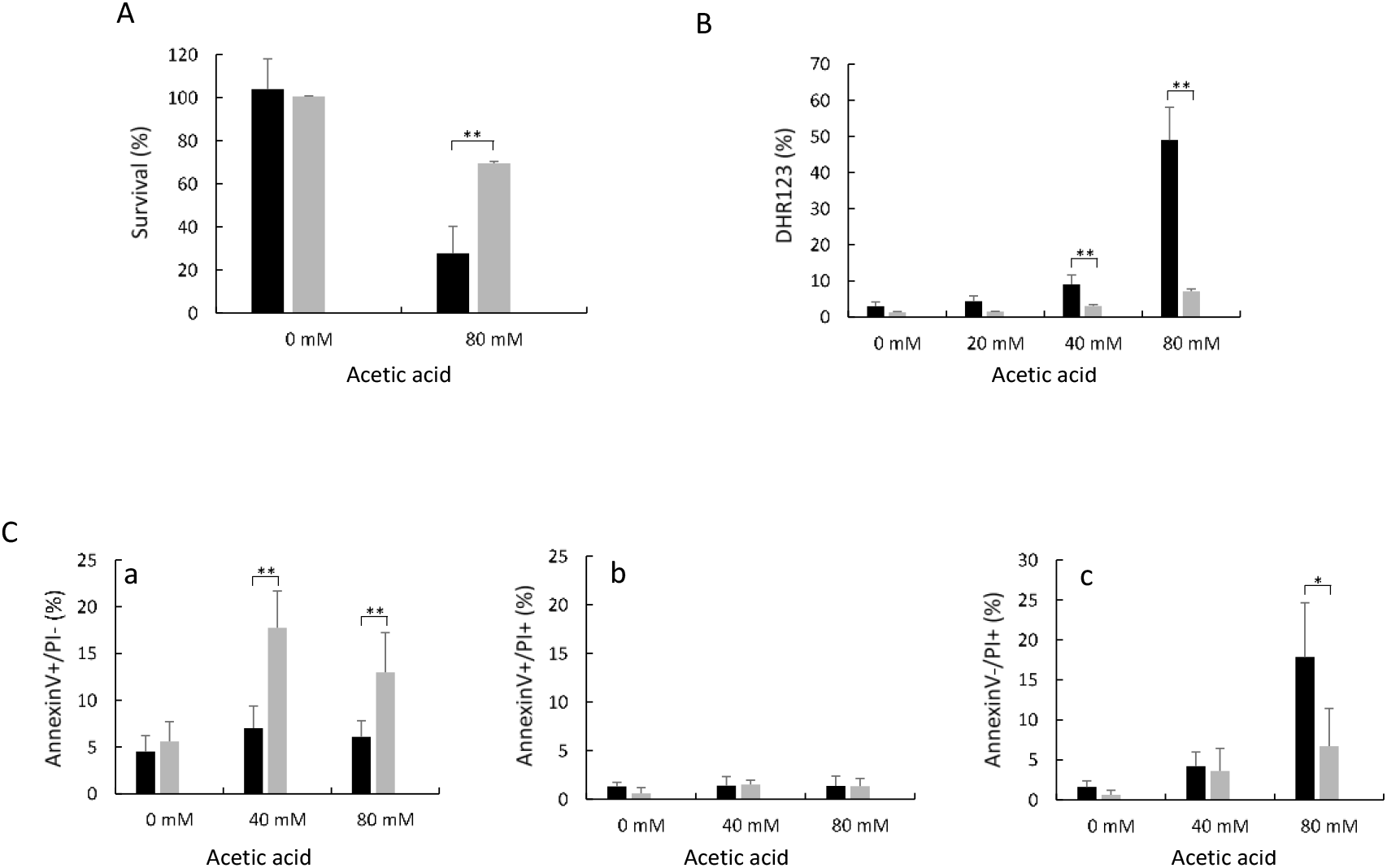
Cell survival, ROS accumulation and cell death in wild-type and ras2Δ cells upon acetic acid treatment. (A) Cell survival of W303-1A and W303-1A *ras2*Δ cells. Cell viability of W303-1A (black bars) and W303-1A *ras2*Δ (grey bars) untreated cells or treated with 80 mM acetic acid was analyzed by measuring colony-forming units (cfu) after 3 days of growth at 30°C. Cell survival is expressed as % to the cfu at time zero. (B) ROS accumulation in W303-1A and W303-1A *ras2*Δ cells. W303-1A (black bars) and W303-1A *ras2*Δ (grey bars) exponentially growing cells were treated with either 20, 40 or 80 mM acetic acid for 200 minutes at 30°C with shaking (160rpm). Dihydrorhodamine 123 (DHR123) was used to assay ROS accumulation. The means of three independent experiments with standard deviations are reported. Student’s t-test *P < 0.05 and **P < 0.01. (C) Assessment of cell death by FITC-coupled annexinV and PI staining. W303-1A (black bars) and W303-1A *ras2*Δ (grey bars) cells growing exponentially on glucose medium were treated with 40 or 80 mM acetic acid for 200 minutes at 30°C, before being processed for determination of phosphatidylserine externalization and membrane integrity by flow cytometry. 30000 events have been evaluated. The means of 3 independent experiments with standard deviations are reported. Student’s-test \**P* < 0.05 and \*\**P* < 0.01.

### 3.3. Active mitochondrial Ras2 protein promote apoptosis and cell death through the cAMP/PKA pathway

Data in literature show that some components of the yeast cAMP/PKA pathway, such as Ira2, adenylate cyclase (Cyr1) [27], and catalytic (Tpk1) and regulatory (Bcy1) subunits of Protein Kinase A (PKA) [28], are also significantly present on mitochondria. Consequently, we investigated whether active mitochondrial Ras2 proteins promote apoptosis and cell death in a cAMP/PKA pathway-dependent manner, since it is well known that Ras proteins in budding yeast activate adenylate cyclase and PKA [15,18,34]. To this aim, we measured cell survival and the levels of ROS and cytological markers of apoptosis and necrosis, in the presence and absence of acid acetic, in two yeast mutants that showed a downregulation of the pathway and reduced activation of PKA: the *gpa2*Δ and *cyr1*Δ strains. In yeast, Gpa2 is a G-protein that activates adenylate cyclase in response to glucose [48]. In the *gpa*2 mutant the cAMP/PKA pathway is downregulated and active Ras proteins are solely localized in the mitochondria [24], while in the *cyr1* strain adenylate cyclase is lacking [35], the cAMP/PKA pathway is totally compromised and a consistent percentage (about 30%) of active Ras proteins is localized in these cell organelles [24]. After treatment with 80 mM acetic acid for 200 minutes at 30°C, both *gpa2*Δ and *cyr1*Δ cells showed a significant increase in cell survival (around 90%) when compared with wild-type cells (Figure 5A). Congruently, the percentage of ROS was significantly reduced in these mutants compared to wild-type cells (Figure 5B). Finally, we measured the level of cytological markers of apoptosis and necrosis in the *gpa2*Δ and *cyr1*Δ mutants. The percentage of *gpa2*Δ and *cyr1*Δ cells in late apoptosis/secondary necrosis following treatment with 80 mM acetic acid was significantly reduced compared to that obtained for the corresponding wild type strain (Figure 4C (c)). As previously shown for the *ras2*Δ strain, also for the *gpa2*Δ and *cyr1*Δ mutants, the percentage of cells in early apoptosis (Annexin V+/PI-), following treatment with 40 mM and 80 mM acetic acid, was significantly high compared with that obtained for the wild strain (Figure 4C (a)). In summary, taken together, our data suggest that mitochondrial active Ras2 proteins promote apoptosis and cell death through the cAMP/PKA pathway.

**Figure 5.**
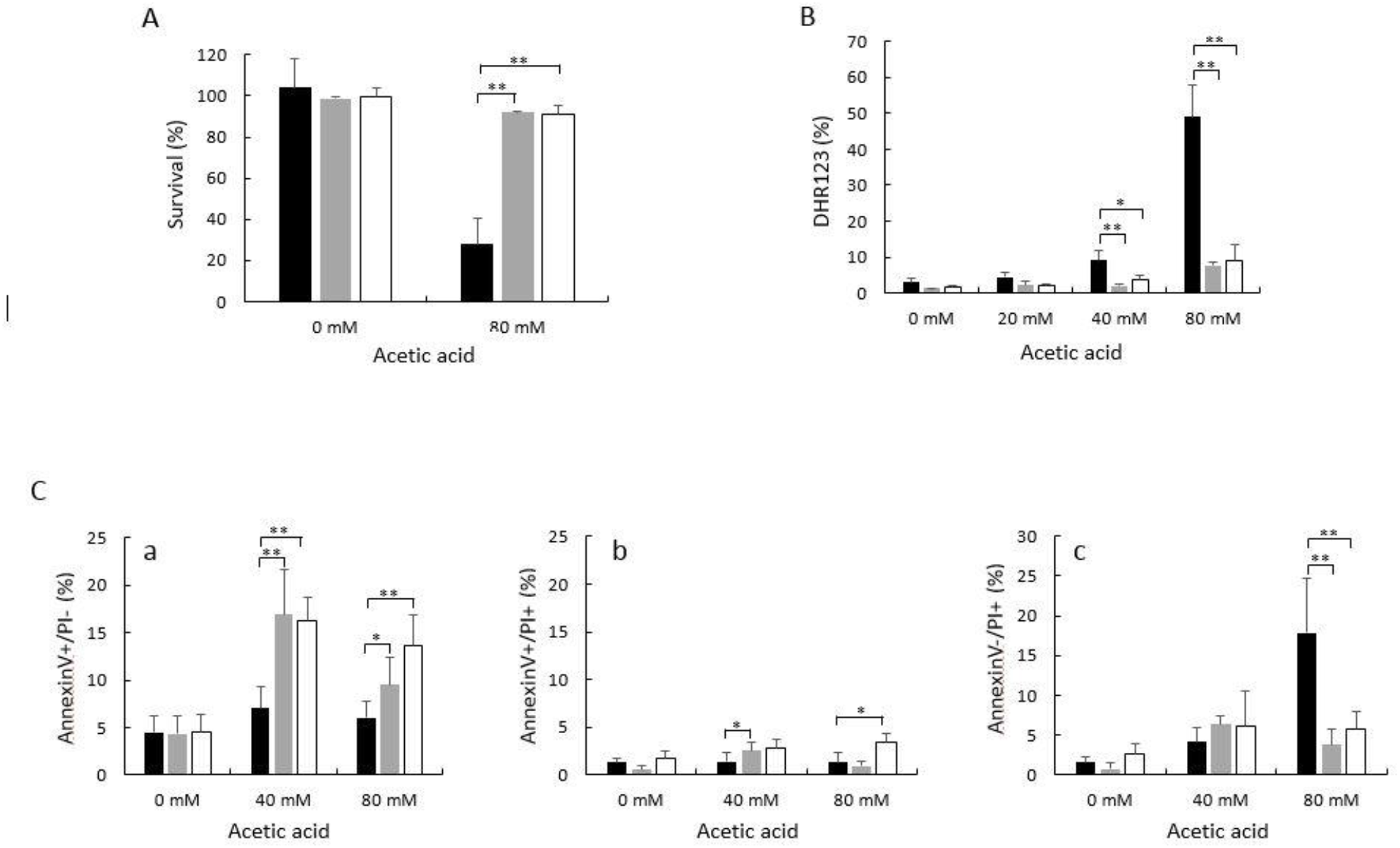
Cell survival, ROS accumulation and cell death in wild-type, *gpa2*Δ and *cyr1*Δ cells upon acetic acid treatment. (A) Cell survival of W303-1A, *gpa2*Δ and *cyr1*Δ strains. Cell viability of W303-1A (black bars), *gpa2*Δ (grey bars) and *cyr1*Δ (white bars) untreated cells or treated with 80 mM acetic acid was analyzed by measuring colony-forming units (cfu) after 3 days of growth at 30°C. Cell survival is expressed as % to the cfu at time zero. (B) ROS accumulation in W303-1A, *gpa2*Δ and *cyr1*Δ cells. W303-1A (black bars), *gpa2*Δ (grey bars) and *cyr1*Δ (white bars) exponentially growing cells were treated with either 20, 40 or 80 mM acetic acid for 200 minutes at 30°C with shaking (160rpm). Dihydrorhodamine 123 (DHR123) was used to assay ROS accumulation. The means of three independent experiments with standard deviations are reported. Student’s t-test *P < 0.05 and **P < 0.01. (C) Assessment of cell death by FITC-coupled annexinV and PI staining. W303-1A (black bars), *gpa2*Δ (grey bars) and *cyr1*Δ (white bars) cells growing exponentially on glucose medium were treated with 40 or 80 mM acetic acid for 200 minutes at 30°C, before being processed for determination of phosphatidylserine externalization and membrane integrity by flow cytometry. 30000 events have been evaluated. The means of 3 independent experiments with standard deviations are reported. Student’s-test \**P* < 0.05 and \*\**P* < 0.01.

## 4. Conclusion

We previously showed that a correlation exists between mitochondrial localization of Ras-GTP and apoptosis. We did that using the eGFP-RBD3 probe, which binds activated Ras proteins with high affinity, but cannot discriminate between Ras1 and Ras2, the two Ras proteins of *S. cerevisiae*. Consequently, in this paper we explored which one of these two GTPases is actually involved in apoptosis and cell death by characterizing the *ras1*Δ and *ras2*Δ mutants concerning localization of active Ras proteins and propensity to undergo these cellular processes, following a pro-apoptotic stimulus. We could show that the Ras2 protein is actually the small G protein, which promotes apoptosis and cell death, and it does that through the cAMP/PKA pathway, while the Ras1 protein might be involved in a pro-survival process. We also demonstrate that lack of *TPS1*, which is known to trigger apoptosis in *S. cerevisiae*, induces localization of active Ras proteins in mitochondria, confirming the correlation between mitochondrial localization of Ras-GTP and apoptosis.

## Acknowledgments

We thank. P. van Dijck, KU Leuven, Belgium, for providing the *tps1* strain, J. Winderickx, KU Leuven, Belgium, for providing the W303-1A *gpa2* strain and J. Thevelein, KU Leuven, Belgium, for providing ST-1 and KP-2 strains. We thank S. Citterio, Università Milano-Bicocca, for her precious technical support and N. Tosetto for technical help.

## Supplementary material

**Supplementary Table I.**
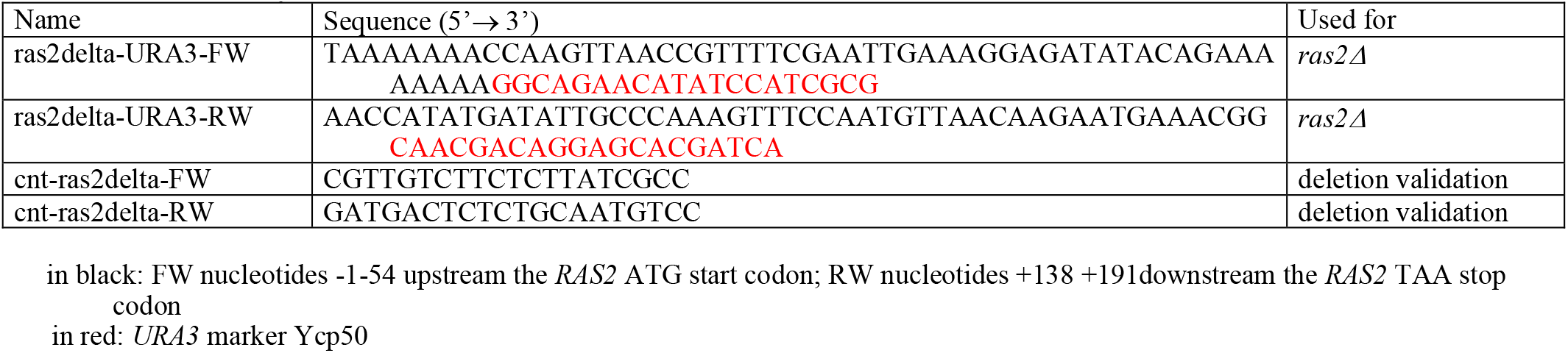
Primers used in this study

**Supplementary Figure 1.**
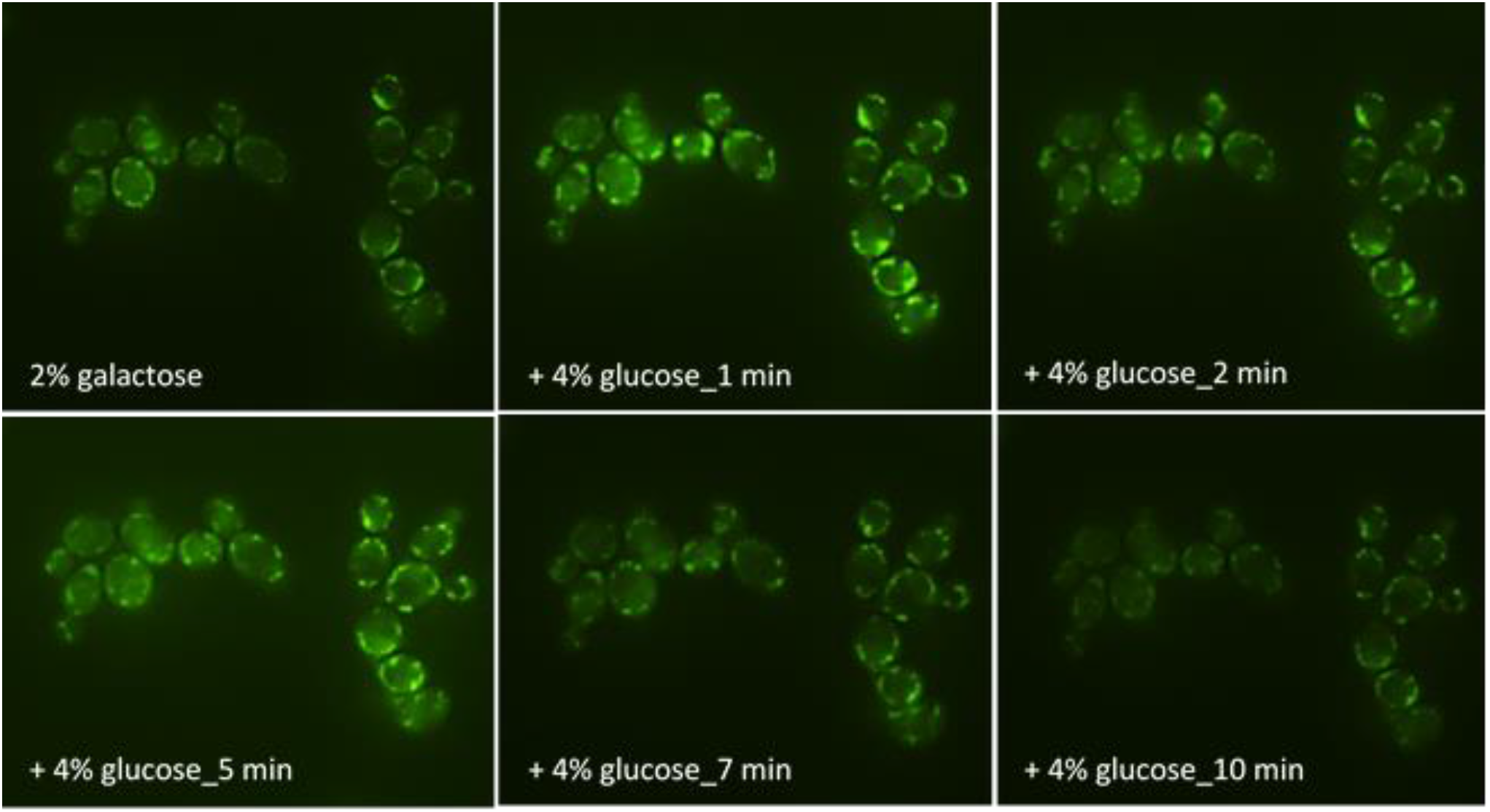
Localization of active Ras proteins in the *tps1*Δ strain before and after addition of glucose to galactose growing cells. Cells expressing the eGFP-RBD3 probe and growing in 2% galactose medium were seeded on concanavalin A-coated cover glass and mounted on top of a Thoma chamber (see Materials and Methods for details). Images were acquired before and after addition of glucose with a Nikon Eclipse 90i microscope equipped with a 60X oil immersion objective and with the same setting of exposure time and imaging (4s, 1.2 gain).

**Supplementary Figure 2.**
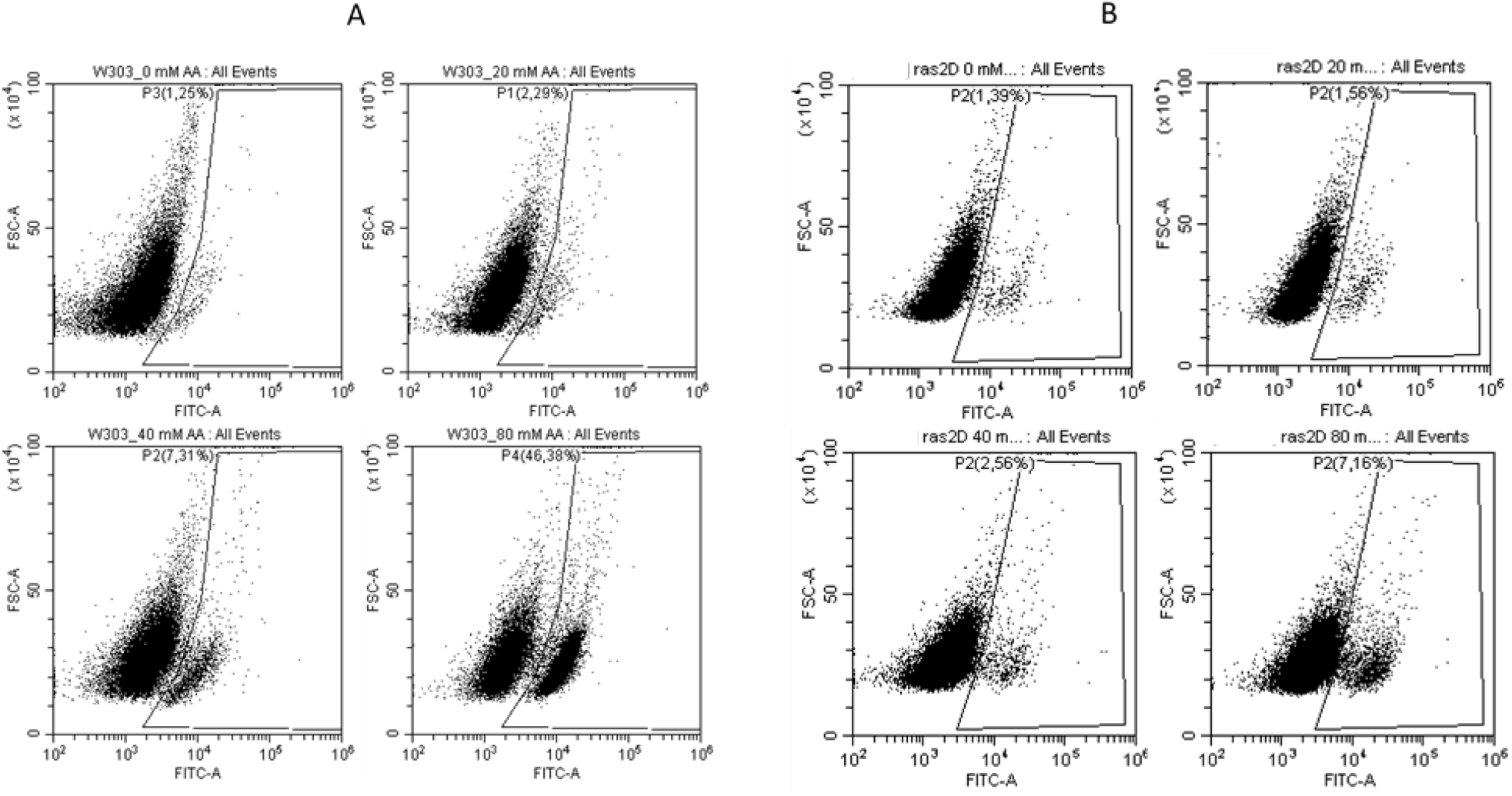
Example of cytofluorimetric analysis after Dihydrorhodamine 123 (DHR123) staining on W303-1A and *ras2*Δ cells. W303-1A (A) and *ras2*Δ (B) exponentially growing cells, untreated or treated with either 20, 40 or 80 mM acetic acid for 200 min at 30° C. On the x-axis the fluorescence (FITC-A) is shown, while on the y-axis the forward scatter (FSC-A) is shown.

**Supplementary Figure 3.**
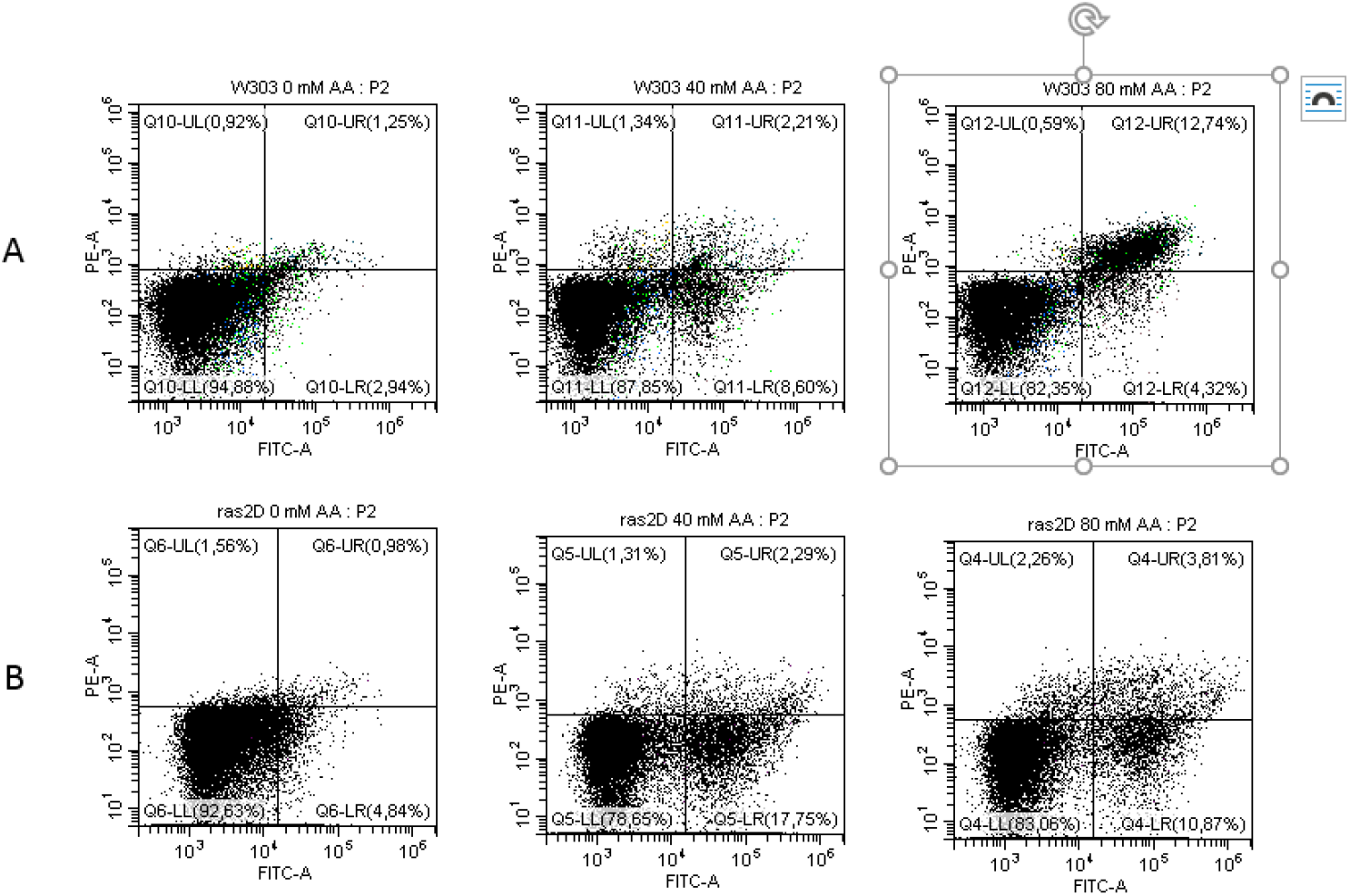
Example of cytofluorimetric analysis after FITC-coupled annexin V and PI staining on W303-1A and *ras2*Δ cells. W303-1A (A) and *ras2*Δ (B) cells growing exponentially on glucose medium were treated with either 40 or 80 mM acetic acid for 200 min at 30° C, before being processed for determination of phosphatidylserine externalization and membrane integrity. Cells in the lower left (LL) quadrant of each cell distribution are annexinV-/PI– and represent the live population. Cells in the lower right (LR) quadrant are annexinV+/PI– and represent the early apoptotic population. Cells in the upper right (UR) quadrant are annexinV+/PI+ and represent the late apoptotic/secondary necrotic population. Cells in the upper left (UL) quadrant are annexinV-/PI+ and represent the primary necrotic population.

